# Comparative transcriptomics shows the evolutionary origin of acoustic fat bodies through a trade-off with skull muscles in toothed whales

**DOI:** 10.1101/2023.05.25.542234

**Authors:** Hayate Takeuchi, Takashi Fritz Matsuishi, Takashi Hayakawa

## Abstract

Toothed whales (odontocetes) have developed acoustic fat bodies, which include forehead melon, extra-mandibular fat body and intra-mandibular fat body. These contribute to the sound transmission pathways in the unique echolocation process of toothed whales. The acoustic fat bodies of Odontocetes accumulate unusual fatty acids, making them novel traits with unique functional anatomy and chemistry. This study provides new insights into the evolutionary origin and lipid utilisation of acoustic fat bodies in harbour porpoise (*Phocoena phocoena*) and Pacific white-sided dolphin (*Lagenorhynchus obliquidens*) using comparative transcriptomics. Comprehensive gene expression analyses indicated that the acoustic fat bodies of toothed whales are composites of muscle and adipose tissues, mainly composed of intramuscular fat. The extra-mandibular fat body is a homologue of the masseter muscle. Additionally, we demonstrated that gene expression-based functional enrichments for specialised branched-chain amino acid metabolism, endocytosis, lysosomes and peroxisomes contribute to the maintenance of the unique fatty characteristics of the acoustic fat bodies of toothed whales. A trade-off occurred in the evolution of acoustic fat bodies as a result of the fatty reorganisation of head muscles involved in aquatic adaptation in toothed whales.

## Background

Understanding the origin of evolutionary novelty is a perennial biological challenge [1,2]. The evolutionary morphological novelty has been defined as structural innovations, such as the development of novel body parts with novel characteristics [1–3]. Novel characteristics often arise from special homologues including the lungs and swim bladders of terrestrial vertebrates and teleosts, respectively [4], and the fins and limbs of fish and tetrapods [5]. Serial homologues that share a subset of a highly conserved gene regulatory network [6–8] have also been identified, including tetrapod forelimbs and hindlimbs [9], and insect head appendages and legs [10].

Toothed whales (odontocetes: suborder Odontoceti), which include porpoises, dolphins and some whales, have echolocation abilities that are lineage-specific. The auditory orientation utilised by toothed whales for underwater navigation, hunting and communication is based on high-frequency clicks. Acoustic fat bodies (AFs), which are specific adipose tissues in the heads of toothed whales, serve as an acoustic transmission pathway for ultrasonic signals [e.g. 11]. AFs are composed of three anatomically distinct tissues: the melon in the forehead, the extra-mandibular fat body (EMFB) outside the mandible and the intra-mandibular fat body (IMFB) in the mandibular foramen (Figure 1). The melon transmits the high-frequency sound produced by the whales’ heads to the water. In contrast, EMFB and IMFB receive sound waves reflected from underwater objects and transmit them to the auditory system [11,12]. The evolution of AFs is viewed as the result of toothed whales’ adaptive evolution to advance in the column and depth of all the world’s oceans.

**Figure 1.**
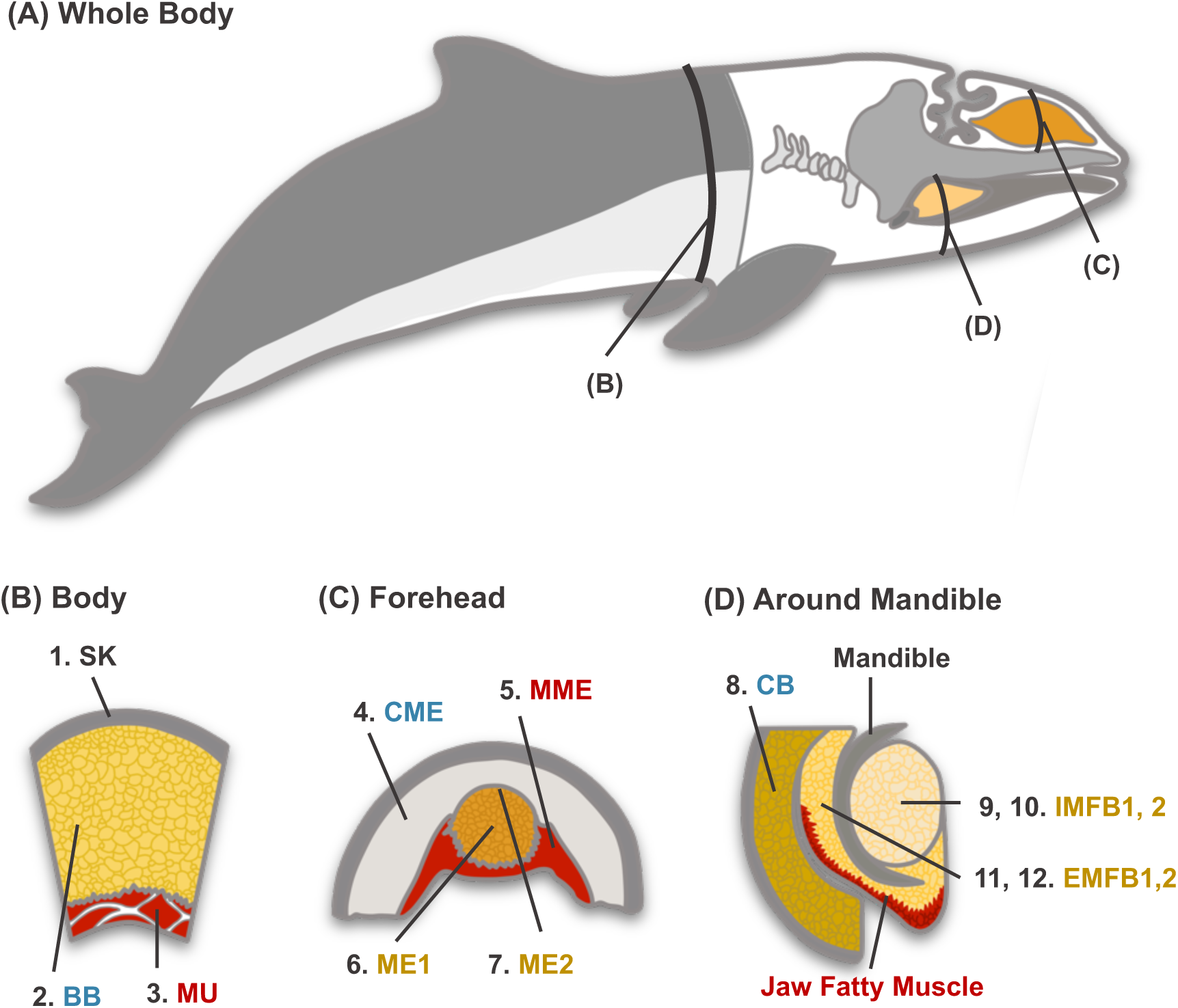
Schematic diagram of acoustic fat bodies and sampling sites. (A) Whole-body planes and (B–D) Coronal planes of sampling sites. Each symbol of the sampling site means as follows: Skin (SK), Body blubber (BB), Muscle (MU), Connective tissue around the melon (CME), Muscle around the melon (MME), Central portion of the melon (ME1), External portion of the melon (ME2), Cranial blubber (CB), Intra-mandibular fat body near the ear bone (IMFB1), Intra-mandibular fat body of the rostral side (IMFB2), Extra-mandibular fat body near the ear bone (EMFB1) and Extra-mandibular fat body of the rostral side (EMFB2). Sampling tissues are classified as ME1, ME2, EMFB1, EMFB2, IMFB1 and IMFB2 as acoustic fat bodies (yellow), MU, MME as comparison tissue of muscle tissue (red), BB, CME and CB as adipose tissue (blue) and SK as outer group tissue (black).

AFs of toothed whales are distinguished by the accumulation of a unique short-branched-chain fatty acid (BCFA) that varies by lineage [13]. Isovaleric acid (*i*-5:0), a BCFA with an odd chain, is abundant in the AFs of delphinoid and phocoenid whales [13–17]. Due to its toxicity when circulating in the bloodstream, no mammals besides humans with a genetic defect are known to accumulate isovaleric acid [18]. The accumulation of isovaleric acid in AFs may result from the endogenous catabolism of branched-chain amino acids (BCAAs; i.e. valine, leucine, isoleucine). Their respective dehydrogenases simultaneously catabolise and convert these three BCAAs to acetyl-CoA or propionyl-CoA. It is hypothesised that the decrease in isovaleryl-CoA dehydrogenase (*IVD*) activity will interfere with the catabolic pathway, resulting in an abnormal accumulation of isovaleric acid (Figure 4C) [13].

Although the evolution of AFs has been extensively studied in the fields of anatomy and lipid chemistry, their genetic basis remains unknown. Identifying homologues of AFs and elucidating the evolutionary origin of unusual fatty acid storage is essential to deciphering the evolution of aquatic adaptation in cetaceans. In this study, we hypothesised that the elucidation of fatty acid utilisation and other identified genetic pathways in toothed whale AF would provide insight into the origin of novelty-AFs. We compared the transcriptomes of AFs and related tissues from two toothed whale species: a harbour porpoise (*Phocoena phocoena*) and a Pacific white-sided dolphin (*Lagenorhynchus obliquiden*s).

During a preliminary anatomical and literature review to determine sampling sites, we identified several AFs of note. First, as is well known, the melon is situated adjacent to the muscle (MME) and the adipose tissue (CME) (Figure 1), but it is more closely associated with the MME than with the CME. Additionally, the surface of the EMFB was densely intertwined with muscle tissue (Figure 1: Jaw fatty muscle) and resembled a masseter muscle. In the melon and extra-mandibular fat body, histological studies have revealed the presence of a distinct muscular component embedded in the adipose mass [19,20]. Furthermore, the IMFB developed adipose tissue within the mandibular foramen, which is typically occupied by the bone marrow. White adipose tissue comprises subcutaneous (SAT), visceral (VAT) and intramuscular (IMAT). Numerous studies and reviews have focused on the distinct metabolic functions of the principal white adipose tissue [21–23]. Recent interest has been drawn to bone marrow adipose tissue (BMAT), which is distinct from white adipose tissue, but research on BMAT is limited[24–27].

Based on these findings, we hypothesised that

(1) AFs are composed of muscle and adipose tissue.
(2) AFs are hypertrophied adipocytes found in muscles (IMAT), with EMFB being homologous to the masseter muscle.
(3) IMFB is also bone marrow-derived adipose tissue (BMAT).

## Materials and methods

### Samples, RNA extraction and sequencing

The efforts of the Stranding Network Hokkaido (SNH) survey resulted in the collection of 12 tissues from a very fresh harbour porpoise (*Phocoena phocoena*; SNH21015) and a Pacific white-sided dolphin (*Lagenorhynchus obliquidens*; SNH21018) [28]. The harbour porpoise used in this study was a bycatch mortality, whereas the white-sided Pacific dolphin was found stranded alive. Both were sampled within hours of death being confirmed. The tissue samples were quickly frozen at −80°C until RNA extraction. Six acoustic fat bodies were obtained: the central portion of the melon, the external portion of the melon (ME1, ME2), the intra and extra-mandibular fat body near the ear (IMFB1, EMFB1) and the intra– and extra-mandibular fat body of the rostral side (IMFB2, EMFB2). Each individual provided two muscle tissues (longissimus dorsi muscle, MU; muscle around the melon, MME) and three adipose tissues (body blubber, BB; cranial blubber, CB; connective tissue around the melon, CME) as well as skin (SK) as an outgroup tissue (Figure 1). When the thickness of the BB along the coronal plane of the dolphin’s axilla was measured, samples of skin, MU and BB were taken from the dorsal side.

Total RNA was extracted using the RNeasy Lipid Tissue Mini Kit (QIAGEN, GmbH, Hilden, Germany), which was designed for optimal lysis of fatty tissues. The purity of the RNA solutions was measured using a NanoDrop 2000 (Thermo Scientific, Waltham, MA, USA). Next, using the Agilent 2200 TapeStation system, the fragment size distribution, the RNA integrity number and the rRNA ratio were estimated (Agilent Technologies). Next, using TruSeq-stranded mRNA, sequencing libraries were constructed (Illumina, San Diego, CA, USA). Additionally, we performed 150-bp paired-end sequencing of these mRNA libraries (RNA-seq) using Illumina NovaSeq6000 (Illumina).

### De novo assembly and functional annotation

Before *de novo* assembly, RNA-seq results (transcriptome) were evaluated for quality using fastp v0.23.2 [29]. Filtering data were obtained by removing low-quality reads, that is, mean phred quality scores below 20, read adapters, too short reads of less than 15 bases, low-complexity consecutive bases in the 3′-end poly-A/T tail and poly-G sequences caused by sequencing artefacts in Illumina NovaSeq series, which are based on their two-colour chemistry. After *in silico* read normalisation, filtered reads for each sample were assembled using Trinity v2.13.2 with the default parameter [30,31]. The EnTAP pipeline was used to annotate assembled contigs [32]. Subsequently, we performed frame selection with TransDecoder v5.5.0 (https://github.com/TransDecoder), functional annotation via similarity search with DIAMOND v0.9.14.115 [33,34] and orthologous group assignment with EggNOG v2.1.10 [35]. The similarity search by DIAMOND was conducted against UniProtKB/Swiss-Prot (release date: February 2022) [36] and NCBI Mammalian RefSeq (ftp://ftp.ncbi.nlm.nih.gov/refseq/release/vertebrate_mammalian/) [36].

The EggNOG databases assigned annotations of gene ontology (GO) and Kyoto Encyclopedia of Genes and Genomes (KEGG) terms [38,39]. In order to remove redundant sequences, CD-HIT v4.8.1 was used to cluster all obtained predicted transcripts for each species at a global sequence identity threshold of 95% [40,41]. On these transcriptome sequences, BUSCO v5.2.2 was used to evaluate the completeness of the assemblies and annotations against the ‘mammalia odb10’ database [42].

### Phylogenetic network analysis

The trimmed reads were mapped to the reference transcriptome using Bowtie 2 v2.4.5 [43] and the abundances of transcript isoforms were calculated using RSEM v1.3.3 [44]. We used the TPM (transcripts per million) matrix to represent relative expression levels that are uniform across libraries [45]. In order to infer phylogenetic network relationships, the TPM values were transformed into binary values (1 or 0) representing the presence or absence of gene expression, with a threshold of TPM = 3. Genes with TPM > 3 were coded as present (1) and those with TPM ≦ 3 were coded as absent (0). This threshold is supported by a statistical model that views the number of transcripts as a mixture of active and inactive expression transcripts [46].

Subsequently, using binary data matrices, the similarity of gene expression profiles between tissues was calculated as pairwise hamming distance, that is, the number of genes whose expression state (1 or 0) differs. A ‘treeness’ of the distance matrix was evaluated by δ statistics [47]. The δ takes values between 0 and 1, which δ approaches a value of 0, indicating complete treeness and δ = 1 indicating a network structure that disrupts treeness. All δ values were calculated for all tetrads (i.e. _12_C_4_ = 495 tetrads in total) from each of the 12 tissues of the harbour porpoise and the Pacific white-sided dolphin. In order to test the hypothesis that AFs are composed of muscle and adipose tissue, we compared the delta values of AF-containing tetrads (i.e. 480 tetrads; δ_AFs_) and AF-free tetrads (i.e. _6_C_4_ = 15 tetrads in total; δ_NAFs_). If AFs were made up of muscle and fat, we would observe large delta values that disrupt the ‘treeness’ between muscle and fat. Next, using in-house Python scripts, 100,000 permutations of bootstrap tests were conducted with δ_diff_ = δ_AFs_ – δ_NAFs_ = 0 as the null hypothesis. Then, using SplitsTree v4.17.1, we performed a phylogenetic network analysis based on the ‘neighbour-net’ method to visualise network relationships between AFs, adipose tissues and muscle tissues [48].

### Differentially expressed genes, specifically expressed genes and enrichment analysis

To extract the genetic characteristics of AFs, we investigated the differentially expressed genes (DEGs) between six AFs (ME1/2, EMFB1/2, IMFB1/2) and three subcutaneous adipose tissues (BB, CB, CME) using the package R DESeq2 [49,50]. We performed a likelihood ratio test with a BH-adjusted *p*-value threshold of 0.05 and a |log_2_(fold change)| >1 as a threshold of DEGs.

We then calculated the ‘Tissue specificity index: τ index’ to identify highly expressed genes that are tissue-specific as follows: [51]. First, highly expressed genes were extracted by removing the gene whose relative expression level was TPM ≦ 3. We then calculated the τ index defined as follows:

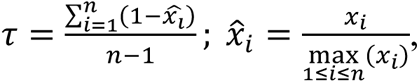

where *x_i_* is the TPM value of the gene in tissue *i* and *n* is the number of tissues. The τ takes values between 0 and 1, in which τ = 1 indicates tissue-specifically expression and τ approaches a value of 0 means ubiquitously expressed in all compared tissues. With > 0.80, acoustic fat bodies-specific highly expressed genes (henceforth AFs-SEGs) were categorised as the most or second-most highly expressed in AFs. Recent comparative research has concluded that this τ index is the best indicator for assessing tissue specificity [52].

In order to evaluate the biological functions of DEGs and SEGs, we performed functional enrichment analysis against the GO and KEGG pathway databases using the clusterProfiler R/BIOCONDUCTOR package [50, 53]. All annotated reference transcripts were used as a background list and a BH-adjusted *p*-value threshold of 0.05 and a *q*-value threshold of 0.05 were chosen as SEG thresholds.

## Results

### Transcriptome assembly and annotation profiling

The Illumina system sequenced a total of 85.8 Gb and 89.3 Gb from 12 tissues of the harbour porpoise and the Pacific white-sided dolphin, respectively (Table 1). The *de novo* assembly of the filtered reads was used to generate the reference transcriptome. The reads assembled reads were combined into a single data set and annotated using the CD-HIT and EnTAP pipelines. The numbers of final annotated transcripts were 69, 569 (porpoise) and 70, 212 (dolphin) transcripts (Table 1). We then evaluated the assembly quality statistics of the reference transcriptome against the complete and fragmented orthologues of highly conserved mammal genes by BUSCO: 72.4% (porpoise) and 72.3% (dolphin) of the orthologues were present as complete and fragmented in the assemblies (Table 2).

**Table 1.** Transcriptome statistics and assembly metrics.

| Species | Tissue | Raw reads | After Trimming reads |  | %Q30 | GC content | Total assembled bases | Contig N50 |
| --- | --- | --- | --- | --- | --- | --- | --- | --- |
|  |  |  | Total reads | Total bases |  |  |  |  |
| Harbour porpoise | SK<sup>a</sup> | 6.656009 G | 43.730337 M | 6.400492 G | 93.257852% | 53.765374% | 670,219,507 | 3,776 |
| Harbour porpoise | MU<sup>b</sup> | 7.420899 G | 48.788474 M | 7.150599 G | 93.714695% | 53.030693% | 344,674,047 | 3,703 |
| Harbour porpoise | MME<sup>c</sup> | 8.002341 G | 52.612422 M | 7.710652 G | 93.760872% | 51.173868% | 548,092,800 | 3,927 |
| Harbour porpoise | BB<sup>d</sup> | 6.154174 G | 40.469958 M | 5.894501 G | 93.505706% | 46.563717% | 211,351,696 | 2,285 |
| Harbour porpoise | CB<sup>e</sup> | 8.534754 G | 56.027510 M | 8.235193 G | 93.623241% | 52.661172% | 1,036,701,453 | 4,065 |
| Harbour porpoise | CME<sup>f</sup> | 6.701294 G | 43.801474 M | 6.433410 G | 93.714939% | 48.279866% | 430,087,813 | 3,037 |
| Harbour porpoise | ME1<sup>g</sup> | 7.364307 G | 48.237312 M | 7.063302 G | 93.815947% | 47.800476% | 511,521,063 | 3,566 |
| Harbour porpoise | ME2<sup>h</sup> | 6.730211 G | 44.150780 M | 6.480375 G | 93.158275% | 49.756490% | 540,886,632 | 3,514 |
| Harbour porpoise | EMFB1<sup>i</sup> | 12.367721 G | 81.019142 M | 11.893544 G | 93.301967% | 51.210294% | 683,063,202 | 3,877 |
| Harbour porpoise | EMFB2<sup>j</sup> | 6.902713 G | 44.792950 M | 6.471157 G | 92.174546% | 49.143097% | 199,058,813 | 1,861 |
| Harbour porpoise | IMFB1<sup>k</sup> | 6.518026 G | 42.703152 M | 6.090932 G | 91.429011% | 49.564174% | 365,749,623 | 3,245 |
| Harbour porpoise | IMFB2<sup>l</sup> | 2.443834 G | 15.613750 M | 2.205062 G | 91.642914% | 52.473959% | 27,018,942 | 430 |
| Pacific white-sided dolphin | SK | 7.516478 G | 49.304040 M | 7.264098 G | 93.099027% | 54.292854% | 943,820,973 | 4,189 |
| Pacific white-sided dolphin | MU | 8.930734 G | 58.702242 M | 8.613530 G | 93.199750% | 48.916646% | 322,088,803 | 3,437 |
| Pacific white-sided dolphin | MME | 6.555734 G | 42.939800 M | 6.303241 G | 93.245243% | 52.302937% | 213,839,825 | 2,692 |
| Pacific white-sided dolphin | BB | 7.487252 G | 49.145166 M | 7.240912 G | 93.881811% | 49.835610% | 714,790,717 | 4,072 |
| Pacific white-sided dolphin | CB | 7.822799 G | 51.481954 M | 7.504348 G | 93.660810% | 53.073039% | 1,048,850,767 | 4,236 |
| Pacific white-sided dolphin | CME | 9.823704 G | 63.938744 M | 9.267085 G | 93.615955% | 54.922172% | 74,118,350 | 1,028 |
| Pacific white-sided dolphin | ME1 | 6.673679 G | 43.816306 M | 6.425037 G | 93.913619% | 47.136894% | 263,248,958 | 2,949 |
| Pacific white-sided dolphin | ME2 | 7.033227 G | 46.079666 M | 6.762142 G | 93.548412% | 50.120124% | 690,973,284 | 4,088 |
| Pacific white-sided dolphin | EMFB1 | 7.230821 G | 47.533394 M | 6.997813 G | 93.719457% | 53.638427% | 385,162,691 | 3,783 |
| Pacific white-sided dolphin | EMFB2 | 7.454592 G | 48.793952 M | 7.199990 G | 92.876471% | 52.638199% | 198,381,499 | 2,567 |
| Pacific white-sided dolphin | IMFB1 | 6.379994 G | 41.637924 M | 6.105715 G | 93.217382% | 52.468262% | 279,025,043 | 3,156 |
| Pacific white-sided dolphin | IMFB2 | 6.424795 G | 41.788552 M | 6.110265 G | 92.319127% | 49.462422% | 361,262,233 | 3,430 |

<sup>a</sup> SK: skin, <sup>b</sup> MU: muscle, <sup>c</sup> MME: muscle around the melon, <sup>d</sup> BB: body blubber, <sup>e</sup> CB: cranial blubber, <sup>f</sup> CME: connective tissue around the melon, <sup>g</sup> ME1: central part of the melon, <sup>h</sup> ME2: external part of the melon, <sup>i</sup> EMFB1: extra-mandibular fat body around the ear, <sup>j</sup> EMFB2: extra-mandibular fat body of the rostral side, <sup>k</sup> IMFB1: intra-mandibular fat body around the ear and <sup>l</sup> IMFB2: intra-mandibular fat body of the rostral side.

**Table 2.** Transcriptome assembly completeness statistics based on BUSCO.

|  | Harbour Porpoise | Pacific white-sided dolphin |
| --- | --- | --- |
| Number of transcripts | 69, 569 | 70,212 |
| Complete BUSCOs | 67.4% (6224)<sup>a</sup> | 67.5% (6236) |
| Single-copy BUSCOs | 46.6% (4301) | 45.2% (4174) |
| Duplicated BUSCOs | 20.8% (1923) | 22.3% (2062) |
| Fragmented BUSCOs | 5.0% (461) | 4.8% (439) |
| Missing BUSCOs | 27.6% (2541) | 27.7% (2551) |

<sup>a</sup> The number in parentheses shows the number of genes.

### Phylogenetic network relationship between acoustic fat bodies, muscles and adipose tissues

Based on statistical data and the neighbour-net method, phylogenetic network analysis revealed intricate network relationships between AFs, adipose tissue and muscle tissue. In order to test the hypothesis that AFs are composites of muscle and adipose tissue, we compared the δ values of AF-containing tetrads with those of AF-free tetrads. Tetrads with AFs had significantly higher δ values than those without AFs for both harbour porpoise and the Pacific white-sided dolphin, as determined by permutation bootstrap tests (*p* < 0.001 in porpoise; *p* < 0.01 in dolphin; Figure 2A, B). The mean and 95% confidence intervals for δ values of tetrads, including AFs in the porpoise and the Pacific white-sided dolphin, were 0.49 [0.47, 0.50] and 0.3 [0.30, 0.33], respectively; δ values of tetrads without AFs were 0.32 [0.25, 0.39] and 0.18 [0.11, 0.25]. This result indicates that AFs distort the treelike structure between muscle and adipose tissue, thereby forming their network relationships. Visualisation of the phylogenetic network using the ‘neighbour-net’ technique also revealed comparable outcomes. The muscle and adipose tissues of the harbour porpoise formed distinct clusters with treelike characteristics (Figure 2C top). The Pacific white-sided dolphin lacked discernible clusters between muscle and adipose tissues (Figure 2D top). In contrast, the phylogeny of the 12 tissues, including AFs, lacked distinct clusters, resulting in network relationships with uncertain treeness (Figure 2C, D bottom). These findings indicate once again that the gene expression profiles of AF are intermediate between those of muscle and adipose tissues.

**Figure 2.**
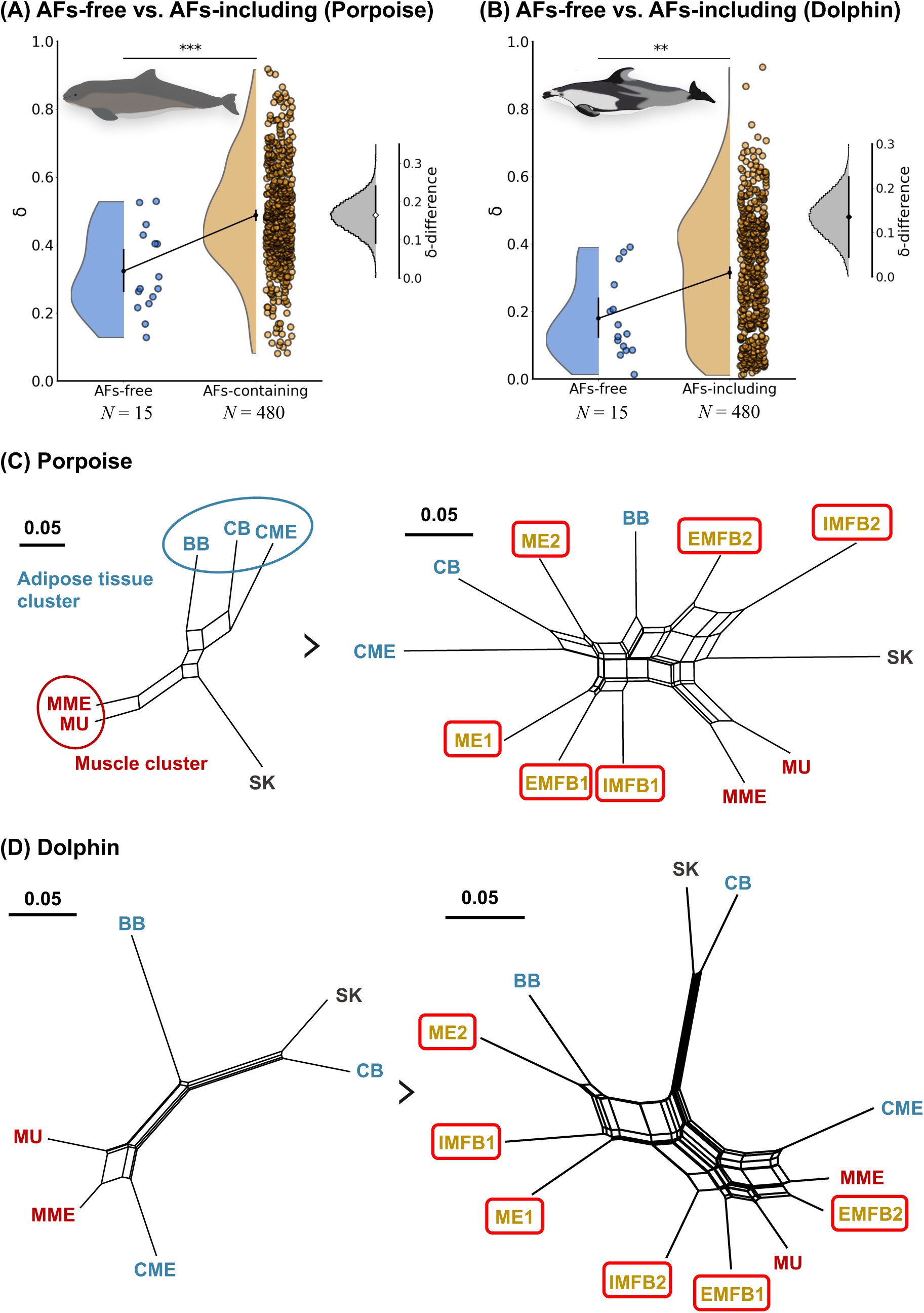
ᵟ-statistics and phylogenetic network relationship of AFs and comparative tissues. (A, B) Violin plots and strip plots of δ value of tetrads including AFs and that of AFs-free, histogram of δ_diff_ obtained by the permutation bootstrap test. δ_diff_ > 0 means that the difference between δ_AFs_ and δ_No-AFs_ is greater than 0. The points and bars in the violin plots or histograms are mean values and 95% confidence intervals of δ or δ_diff_. Figures were drawn using the Python Seaborn library. (C, D) Visualised phylogenetic networks without AFs and all 12 tissues were drawn by SplitTree4.

### Acoustic fat bodies originating from intramuscular adipose tissue

Melon and EMFB have been shown to activate muscle-related genes. In order to deduce the evolutionary origin of AFs from the tissue specificity of gene expression, functional enrichment analysis of specifically expressed genes (SEGs) was performed. Melon– and EMFB-SEGs were significantly enriched for numerous muscle-related KEGG/GO terms, including hsa05412: Arrhythmogenic right ventricular cardiomyopathy and GO:0060537: muscle tissue development in both the harbour porpoise and the Pacific white-sided dolphin (Figure 3A, Table S1–24). EMFBs have muscle-related functions comparable to those of the longissimus dorsi muscle (MU) (Figure 3A, Table S15, 16, 21, 22). In the Pacific white-sided dolphin, AFs, including IMFB, were more enriched for muscle-related KEGG/GO terms than in the porpoise. Given that our preliminary dissections revealed that the mandibular fat bodies of porpoises are fatter than those of other toothed whales, this result is reasonable (data not shown). These findings are consistent with the scattered evidence that AFs are evolutionarily closely related to both adipose tissues and muscles.

**Figure 3.**
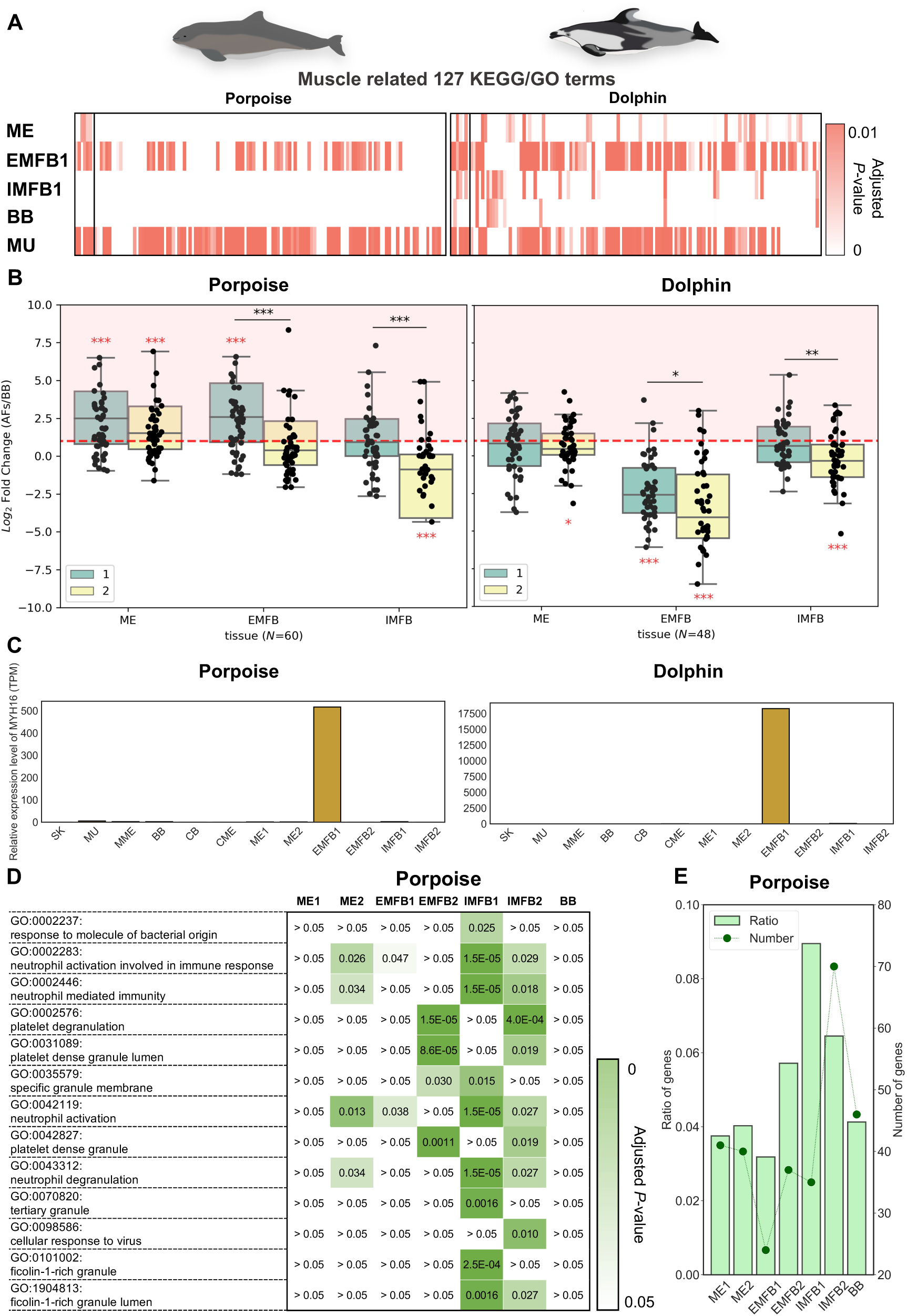
Gene expression analyses imply the evolutionary origin of acoustic fat bodies. (A) Heatmaps of 127 KEGG/GO enrichment terms associated with muscle in acoustic fat bodies (ME: melon, EMFB: extra-mandibular fat body near the ear bone, IMFB: intra-mandibular fat body near the ear bone), muscle (MU) and body blubber (BB). In this graph, the adjusted *P-value* for over-representdness <0.05 (KEGG terms) and < 0.01 (GO terms) are shown. (B) Box plots of log2 fold change of acoustic fat bodies (AFs) compared with body blubber (BB) in previously reported that differentially expressed genes of highly expressed in bovine intramuscular adipose tissue than in subcutaneous fat (Lee et al. 2013). See the text in detals (*: *p* < 0.05, **: *p* < 0.01, ***: *p* < 0.001). (C) Bar plots of relative expression levels (TPM) of *MYH16* in 12 tissues (left: harbour porpoise, right: Pacific white-sided dolphin). (D) A heat map of immune-related terms enriched in AFs by functional enrichment analysis of gene ontology in the harbour porpoise. Adjusted *P-value* for over-representation <0.05 was shown in this graph. (E) Bar plot (ratio) and line plot (number) of the number and proportion of BM-marker genes (see the text in detals) among specifically expressed genes in AF in the harbour porpoise.

We suggested that Melon and EMFB originate from intramuscular adipose tissue (IMAT). It has been reported that genes differentially expressed at higher levels in bovine IMAT than in their subcutaneous fat (SAT) explain the differentiation of these traits (here IMAT-marker genes) [21]. We compared the expression levels of these IMAT-marker genes in AFs versus SAT blubber (BB). The porpoise ME and EMFB expressed IMAT-marker genes at significantly higher levels than BB (Wilcoxon signed-rank test in R software, ME1: *p* < 0.001, ME2: *p* < 0.001, EMFB1: *p* < 0.001; Figure 3B left). No significant increase in IMAT-marker genes expression was detected in the IMFB of the porpoise (Wilcoxon signed-rank test; Figure 3B left). On the other hand, the expression of IMAT-marker genes was not significantly increased in the AFs of the Pacific white-sided dolphin (Wilcoxon signed-rank test; Figure 3B right). The EMFB/IMFB near the ear bone (EMFB1/IMFB1) expressed IMAT-marker genes at a significantly higher level than the EMFB/IMFB of the rostral side (EMFB2/IMFB2) (Brunner-Munzel test in R package ‘lawstat’, EMFB of porpoise: *p* < 0.001, IMFB of porpoise: *p* < 0.001, EMFB of dolphin: *p* < 0.01, IMFB of dolphin: *p* < 0.05; Figure 3B) [54]. Our anatomical examinations demonstrated that EMFB2 has dense muscle and that Pacific white-sided dolphins have muscularity AF (data not shown). As in previous results, the expression pattern of muscle-related genes in AFs may have a significant effect on our findings. These results summarise the IMAT origin of ME and EMFB and support our hypothesis that ME and EMFB are caused by the hypertrophy of developed fat masses in muscle tissue.

In addition, we discovered that EMFB is an evolutionary homologue of the masseter muscle. Myosin heavy chain 16 (*MYH16*) is a member of the myosin gene family, one of the main proteins that make up muscle myofibrils and a type of molecular motor expressed exclusively in the masticatory muscle of mammals [55]. *MYH16* was only highly expressed in EMFB1 in the harbour porpoise and the Pacific white-sided dolphin (Figure 3C). This is the first report to genetically demonstrate that the EMFB gene of toothed whales is homologous to the masseter muscle gene.

Active immune-related gene function is present in IMFB. Enrichment analysis of IMFB-SEGs revealed that porpoise IMFB was enriched in immune-related functions such as complement and coagulation cascades, GO:0002283 neutrophil activation involved in the immune response, GO:0002446 neutrophil-mediated immunity and GO:0002237 response to the bacterial molecule (Figure 3D, Table S17, 18). In contrast, not only IMFB but also the majority of tissues of the Pacific white-sided dolphin were enriched for infectious disease and cancer, including hsa05169 Epstein-Barr virus infection, hsa05202 transcriptional misregulation in cancer (Table S19–24). The Pacific white-sided dolphin used in the study was pregnant and live-stranded and may have been infected with severe diseases.

Our analysis did not fully support the hypothesis that IMFB originates in the bone marrow. 165 genes have been reported to be highly and specifically expressed in human bone marrow (BM-marker genes) [56]. We determined the number and proportion of BM-marker genes in porpoise AFs-SEGs. The number and proportion of BM-marker genes were marginally higher in IMFB-SEGs than in other AF-SEGs, but no clear trends were observed (Figure 3E). We hypothesised that IMFB is bone marrow-derived adipose tissue (BMAT) because the IMFB typically develops at the site of bone marrow. The distinct bone marrow origin of IMFB was not supported, despite the fact that these results suggested that IMFB may be an immune function-rich tissue.

### Unique lipid metabolisms formed acoustic fat bodies

As anticipated, it was proposed that the accumulation of isovaleric acid (*i*-5:0) in AFs is the result of a modification of the general leucine degradation pathway. To rationalise the different lipid metabolism pathways that form AFs, differentially expressed genes were identified between six AFs (ME1/2, EMFB1/2, IMFB1/2) and three adipose tissues (BB, CB, CME) and functional enrichment analysis was conducted. The genetic function of extracellular matrix (e.g. GO: 0030198 extracellular matrix), branched amino acid degradation (e.g. hsa00280 valine, leucine and isoleucine degradation), endocytosis (e.g. GO: 0045807 positive regulation of endocytosis), lysosome (e.g. GO: 0043202 lysosomal lumen) and peroxisome (e.g. GO: 0005777 peroxisome) were significantly enriched (Figure 4A, Table S25–28). BCAAs are normally degraded in the liver, but free BCAAs are metabolised as energy sources in the muscle [57]. Genes involved in the common degradation pathway of all BCAAs (*BCAT1*, *BCAT2*, *BCKDHA*, *BCKDHB* and *DBT*) were expressed at significantly higher levels than in the longissimus dorsi muscle (MU) (Figure 4B, C, Table S29). Conversely, as expected, the expression level of *IVD*, which catalyses the conversion of isovaleryl-CoA to acetyl-CoA, decreased in AFs relative to MU (Figure 4B, C, Table S29). These findings support Koopman’s (2018) hypothesis that the *i*-5:0 that accumulates in the AFs of toothed whales is synthesised from leucine.

**Figure 4.**
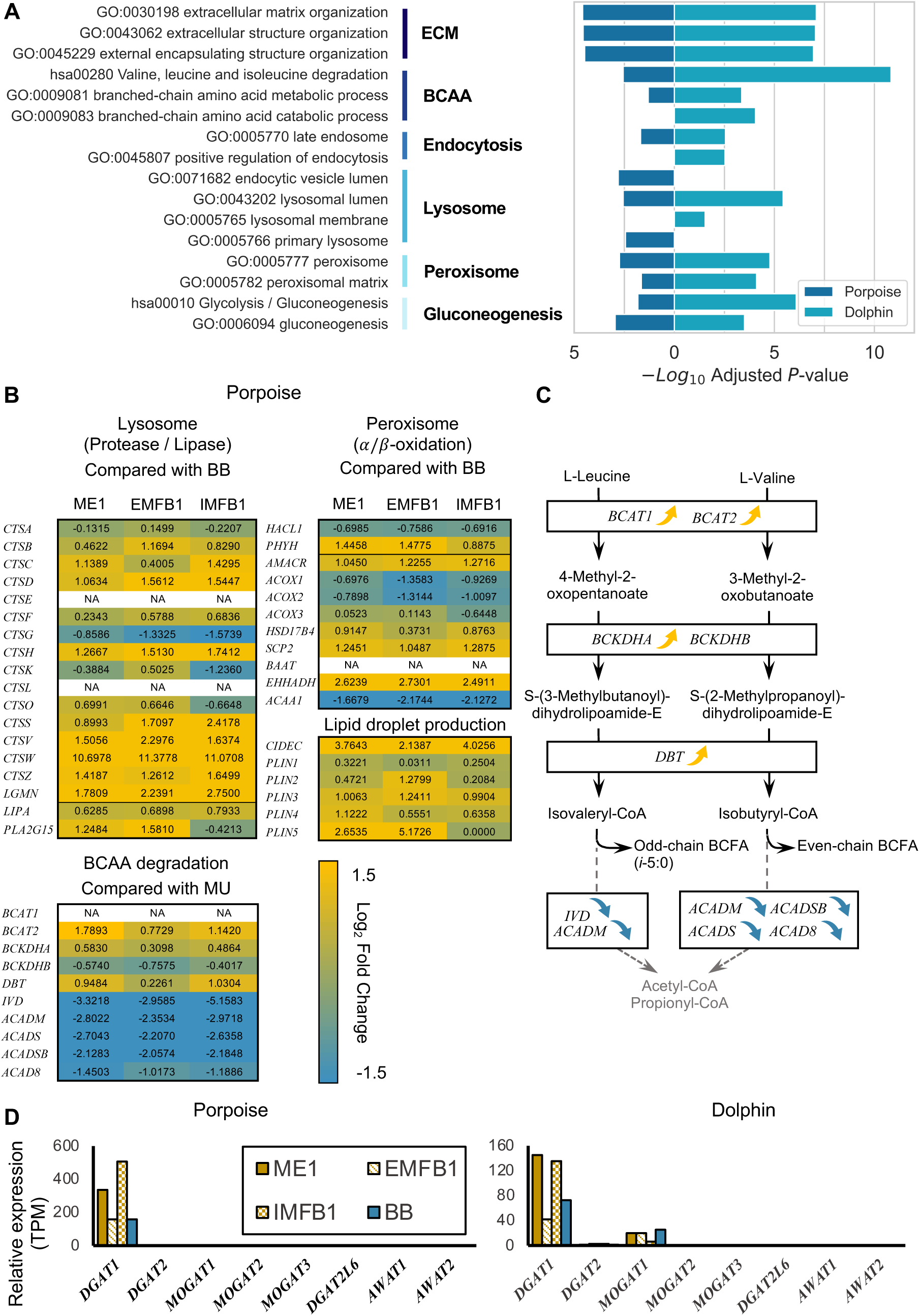
KEGG/GO enrichment analysis and Relative expression profile (TPM) of genes involved in unique use of lipids in acoustic fat bodies (AFs). (A) Functional enrichment analysis of genes differentially expressed higher in AFs than in adipose tissue (BB: body blubber, CB: cranial blubber, CME: connective tissue around the melon) in the harbour porpoise (left) and the Pacific white-sided dolphin (right). (B) Heatmaps of the log2-fold change of genes involved in lysosomes and peroxisomes comparing TPM between AF and body blubber in the harbour porpoise. A Heatmap of the Log_2_ fold change of genes involved in branched-chain amino acid degradation comparing TPM between AFs and muscle in the harbour porpoise. NA means genes that are not annotated. (C) Expression trends of genes associated with the branched-chain amino acid-branched-chain fatty acid synthesis pathway in harbour porpoise. (D) Bar plots of the relative expression level (TPM) of the *DGAT1* family and *DGAT2* in the 12 tissues of the harbour porpoise (left) and the Pacific white-sided dolphin (right).

The odd-chain BCFA-biased characteristics of AFs cannot be explained solely by evolution via modification of the BCAA degradation pathway. Contrary to expectations, we demonstrated that AF decreased the expression of genes (*ACADM*, *ACADSB*, *ACADS* and *ACAD8*) that catalyse the conversion of isobutyryl-CoA to propionyl-CoA (Figure 4B, C, Table S29). Not only the leucine pathway but also the valine pathway was altered, indicating an evolutionary consequence of the BCAA degradation pathway that enabled the potential production of even-chain BCFA.

The functions of lysosomes and peroxisomes were prominent in harbour porpoise AFs. Lysosomes serve primarily to degrade and recycle macromolecules. In the KEGG pathway 04146: Lysosome, the expression levels of proteases (*CTSA* family) and lipases (*LIPA*, *PLA2G15*) functioning in lysosomes were higher in AFs than in BB (Figure 4B, Table S29). Peroxisomes are highly conserved in eukaryotes and have a distinct fatty acid oxidation system than mitochondria. Beta oxidation degrades the fatty acid chain by two carbons [58] and alpha oxidation shortens it by one carbon each [59,60]. A comparison of expression levels of genes involved in the KEGG pathway (hsa04146: peroxisome) revealed that genes for alpha oxidation (*PHYH*) and beta oxidation (*AMACR*, *HSD17B4*, *SCP2* and *EHHADH*) were more highly expressed in AFs than in BB (Figure 4B, Table S29). Consequently, the adaptive evolution of biological functions other than the BCAA degradation pathway, such as endocytosis, lysosomes and peroxisomes may have contributed to the formation of unbalanced odd-chain BCFA-biased AFs.

In AFs, the conversion of synthesised fatty acids into lipids and their storage as lipid droplets are active. *DGAT1* and *DGAT2* gene familiy (*DGAT2*, *DGAT2L6*, *MOGAT1*, *MOGAT2*, *MOGAT3*, *AWAT1*, *AWAT2*) catalyse the synthesis of lipids from fatty acids [61–63]. Both the porpoise and the Pacific white-sided dolphin exhibited elevated DGAT1 activity (Figure 4D). Furthermore, the expression of *MOGAT1* was detected in the Pacific white-sided dolphin. *CIDEC* and *PLIN1/2/3/4/5* are essential for the storage of synthesised lipids as lipid droplets [64–66]. These genes were found to be expressed at a higher level in AFs than in BB in the harbour porpoise (Figure. 4B, Table S29). These findings suggest that the evolution of unique lipid utilisation genetic pathways in AFs that serve as acoustic transduction pathways contributes to the maintenance of stable adipose tissue.

## Discussion

We carried out transcriptome analyses of AFs and comparative tissues of a harbour porpoise and a Pacific white-sided dolphin to determine the origin of AFs and the evolution of their distinctive lipid chemistry. Phylogenetic network analysis using δ statistics and the neighbour-net method revealed that AFs, muscles and adipose tissues have network-like connections (Figure 2). This indicates that AFs are evolutionarily related to both adipose tissues and muscles. Next, comparative gene expression analysis of our data and previous studies revealed that Melon and EMFB are hypertrophic components of adipose mass within the muscle tissues (IMAT) rather than the subcutaneous fat covering the body surface (Figure 3A, B). In addition, we provided evidence for the EMFB-specific expression of *MYH16* (Figure 3C), proving that EMFB is a special homolog of masticatory muscle. Furthermore, our meticulous genetic pathway analysis has shed light on the abnormal fatty acid (*i*-5:0) storage system of AFs. According to our findings, modification of the leucine degradation pathway is required for the synthesis of *i*-5:0 (Figure 4B, C, Table S29). Thus, based on the functional enrichment analysis of AFs, we propose the evolution of a novel *i*-5:0 storage pathway (details below; Figure 4, Figure 5, Table S27, 28).

**Figure 5.**
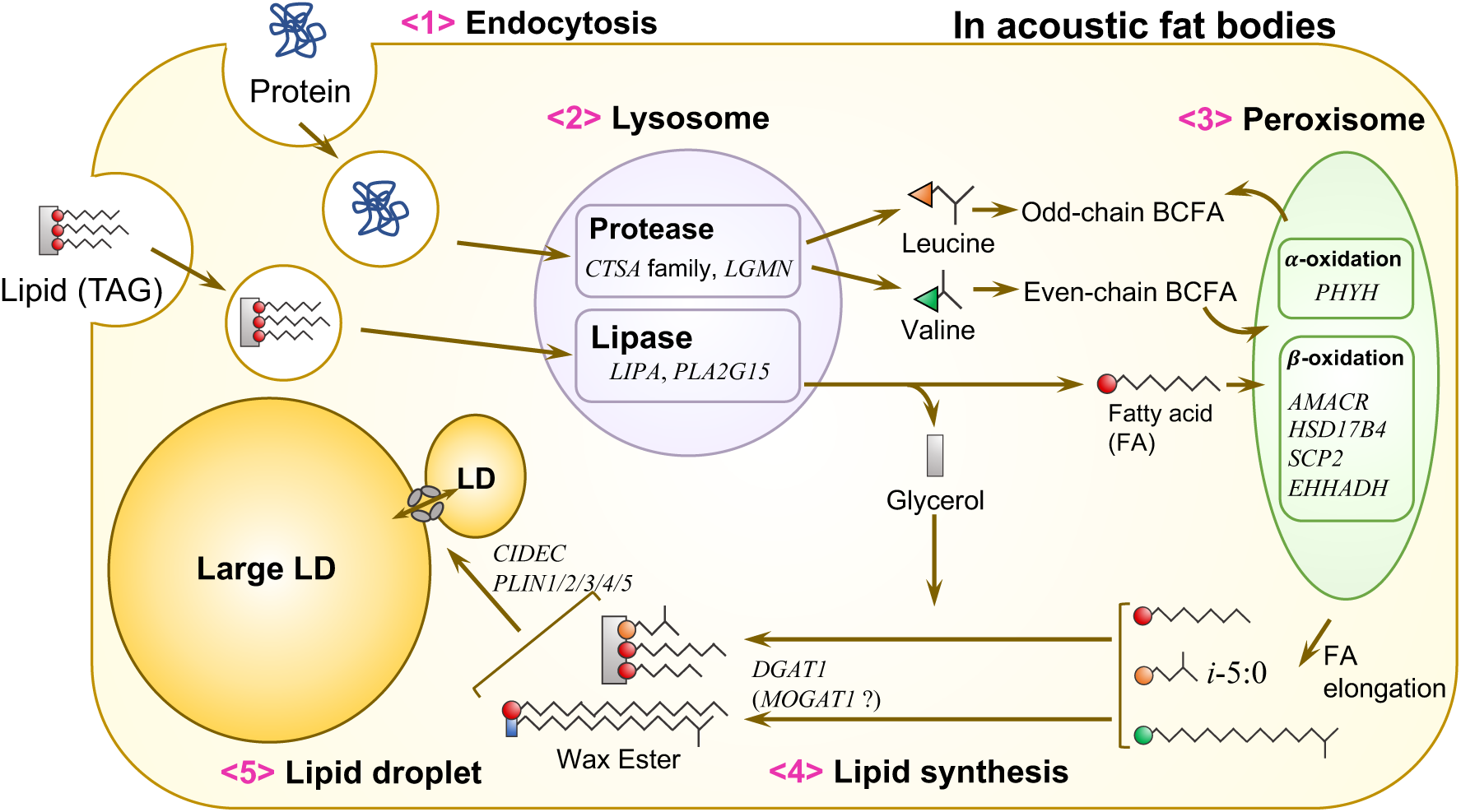
Proposed schematic diagram of genes associated with specialised lipid utilisation pathways acoustic fat bodies of (AFs) of toothed whales. We proposed a genetic pathway to store isovaleric acid (*i-*5:0) built in some toothed whales (see text in details).

### Limitations of this study

It should be noted that the small sample size used in this study may have led to erroneous conclusions. In this investigation, only two-stranded or bycatch individuals’ RNA was sequenced in bulk. Due to the aquatic nature and rarity of wild cetaceans, it is difficult to obtain large quantities of very fresh samples for RNA research. Therefore, this study lacked sufficient samples for statistical analysis. In addition, the Pacific white-sided dolphin that died as a result of beaching was pregnant. According to our genetic analysis, this individual may have suffered from a serious illness, such as an infection. In other words, it is possible that the Pacific white-sided dolphins used in this study deviate from the genetic pathway of healthy dolphins.

Instead of hierarchies, we chose to depict the evolutionary phylogeny of AFs using networks. The scRNA-seq-derived delta statistic has a mode near the δ = 0 and has a long tail extending to δ = 1 [67,68]. Although the bulk RNA-seq in this study, the mode of the δ was between 0.4 and 0.5, the frequency distribution of δ in the harbour porpoise has a long tail that extends to both δ = 0 and δ = 1. Our analysis validated the representation of bulk sequencing data at the tissue level as a network relationship rather than hierarchical clustering (Figure 2). Bulk sequencing data, however, are significantly affected by abundant cells in the sample. Additionally, adipocytes are known to store high levels of lipids, resulting in fewer cells per sampling volume and lower RNA extraction yields [69]. Therefore, it was necessary to account for the possibility that the myocytes slightly contained in AFs skewed our results. Some results of the Pacific white-sided dolphin, whose AFs are muscular, were more difficult to interpret than those of the porpoise (Figure 2D, Figure 3B). In order to overcome these limitations, researchers will need to continue collecting samples and addressing the scRNA sequencing at the cellular level.

### Melon originating from intramuscular adipose tissue

Our findings indicated that the melon descend from IMAT. Activated ECM-related genes are present in AFs (Figure 4A, Table S27, 28). ECM is essential for adipocyte growth and differentiation [70]. Significant differences in ECM-associated gene expressions between IMAT and SAT have been reported [21]. We demonstrated that these genes are expressed at a higher level in melon than in BB (Figure 3B).

ME-SEGs were also enriched for cardiomyopathy-related function, such as hsa04510 arrhythmogenic right ventricular cardiomyopathy, through KEGG functional enrichment analysis. Notably, arrhythmogenic right ventricular cardiomyopathy is known to result in myocardial repair through fibrofatty replacement (information from KEGG pathway: hsa05412). The melon of the bottlenose dolphin is tightly coupled to the facial muscles, indicating that it may modify the beam width and directionality of acoustic beams emitted and passing through the melon [71]. Combining our genetic-level analysis, anatomical data and echolocation clicks through melon tuned by muscle, it is plausible that melon originates from IMAT hypertrophy in muscle tissue.

### EMFB, the special homologue of the masticatory muscle

Our discovery of EMFB1-specific *MYH16* expression demonstrates that EMFB is a special masseter muscle homologue (Figure 3C). As *MYH16* is known to be expressed in masticatory muscles, particularly in the tensor veli palatini muscle, which plays a role in opening the auditory tube and is located near the ear [55, 72]. Therefore, it appears that *MYH16* expression was not detected in the rostral region of EMFB (EMFB2). In the context of their phenomenal aquatic adaptations, cetaceans have significantly redesigned their auditory system and adapted to a feeding ecology in which they swallow food without chewing. Thus, the muscle strength of the tensor veli palatini muscle became an unnecessary relic in toothed whales, resulting in the formation of an unprecedented EMFB by fatty replacement of the masseter muscle.

### IMFB specialising in immune system function

In this study, we were unable to fully determine the evolutionary origin of IMFB. Although IMFB appeared to have excellent immune-related functions (Figure 3D), no broad association with bone marrow, the origin of immune systems, was identified (Figure 3E). Notably, the rostral side of IMFB (IMFB2) did not appear to be related to IMAT (Figure 3B). Since the IMFB is connected to the EMFB on the side of the ear bone (Figure 1D), IMFB1 may contain IMAT. In contrast, IMFB2 is deeply buried in the mandibular foramen and is distant from EMFB (data not shown).

However, some researchers believe that IMFB originated in the bone marrow [73,74]. Some components of the venous circulation in IMFB are comparable to the ‘open’ circulation in the bone marrow [73]. Our comprehensive literature search at the single-gene level also suggests IMFB’s myeloid relevance. Some genes with significant associations with bone marrow function were identified in IMFB-SEGs shared by the harbour porpoise and the Pacific white-sided dolphin. The matrix Gla protein (*MGP*) inhibits extracellular matrix ossification in the bone marrow and is essential for adipocyte differentiation in the bone marrow [75]. It has also been demonstrated that *DLX5* overexpression inhibits osteogenic differentiation of bone marrow stromal cells [76]. Furthermore, *RIN3* negatively regulates the bone formation, as RIN3-knockout mice exhibited an increase in trabecular bone [77]. The bone marrow haematopoietic stem cell (HSC) differentiation is promoted by silencing endogenous G0S2 expression, supporting an inhibitory role for *G0S2* in HSC proliferation [78]. We cannot rule out the possibility that IMFB is the developed adipose tissue as a result of suppressing ossification and HSC differentiation in the bone marrow in light of these credible previous studies.

### Evolution of genetic pathways behind odd-chain BCFA dominated AFs

Our exhaustive gene expression experiments demonstrated that *i*-5:0 in AFs is derived from the leucine residue (Figure 4B, C, Table S29). Several previous studies have suggested that modification of the leucine degradation pathway alone, without alteration of the valine pathway, generates AFs with a high concentration of *i*-5:0 [e.g. 13]. However, we provided evolutionary evidence of modified gene pathways for the degradation of both leucine and valine. Such genetic pathway modifications enable the potential synthesis of even-chain BCFAs in AFs, which is inconsistent with *i-*5:0-dominated AFs.

Our gene function enrichment analysis suggests a different metabolic pathway for the *i*-5:0 storage in AFs (Figure 5). First, AFs acquire extracellular proteins through endocytosis. Second, the action of lysosomes degrades proteins into BCAAs and other amino acids. Third, peroxisomal alpha oxidation functions in the synthesis of precursors for odd-chain-biased fatty acids (*i*-5:0). Finally, the synthesised *i*-5:0 is converted to lipids and stored as droplets of lipids. High expression levels of lysosomal proteases in AFs (Figure 4B, Table S29) indicate the significance of the amino acid recycling mechanism by extracellular protein degradation in lysosomes for the recycling of amino acids. The ability to convert even-chain BCFAs into odd-chain BCFAs can be granted by alpha oxidation boosts that shorten carbon chains one by one. Therefore, the formation of AFs with *i-*5:0-biased can be explained by alpha oxidation within AFs.

In the AFs of the Pacific white-sided dolphin, a certain level of lipid synthesis by *MOGAT1* was observed (Figure 4D). Several delphinoid AFs appear to contain specialised lipids (wax esters) [13]. *MOGAT1* may be involved in wax ester synthesis in the dolphin family Dolphinidae’s Pacific white-sided dolphin.

Unlike subcutaneous fat, the AFs of toothed whales are unaffected by starvation or ambient temperature and never become thin. This is because AFs must serve as pathways for echolocation clicks regardless of season or nutritional status. The extraordinary *i*-5:0 storage as lipid droplets allow toothed whales to maintain stable AFs. The formation of AFs from IMAT (or BMAT) distinct from subcutaneous fat, which is fluidly modified by external factors, may have facilitated the evolution of the auditory system of toothed whales.

## Conclusion

Through comparative transcriptomics of acoustic fat bodies and related tissues in a harbour porpoise and a Pacific white-sided dolphin, we hypothesised that elucidation of the evolution of lipid utilisation and other notable gene pathways in toothed whale acoustic fat bodies will provide essential insights into how the novel trait-AFs arose. We demonstrated that the fat of the origin of the melon and mandibular fat body (EMFB) is an enlarged adipose mass (IMAT) within muscle tissue. EMFB is a homologue of the masticatory muscle, in particular. IMFB is specialised for immune system function and may consist of bone marrow-derived adipose tissue (BMAT).

Furthermore, we explained that *i-*5:0-enriched AFs in some toothed whales may have resulted from the evolution of complex cellular functions of endocytosis, lysosomes, and peroxisomes, in addition to the modification of the leucine degradation pathway.

Significant changes in skull morphology and the transition to a feeding ecology that does not require masticatory ability led to a trade-off in the construction of toothed whale AFs. AF may have originated from intramuscular fat (or bone marrow fat) as opposed to subcutaneous fat, which changes lipid stores and shapes in response to external stimuli. The evolution of adipose tissue division of labour has enabled the de novo adoption of AFs as acoustic transmission pathways in the aquatic adaptation of the toothed whales’ auditory system.

## Ethics approval

The authors collected the specimens with the Stranding Network Hokkaido (SNH). The samples were collected in compliance with Japan’s relevant laws and regulations and SNH provided them unconditionally for this study.

## Availability of data and materials

RNA-seq data determined in this study were deposited in the DDBJ database with accession numbers PRJDB14714.

## Author contributions

HT conceived and designed the research. TFM and TH supervised the research. TFM and HT collected genetic samples. HT performed molecular biology experiments. HT performed informatics analyses. All authors interpreted and discussed data. HT wrote the original draft. All authors edited and approved the final version of the manuscript.

## Acknowledgements

We thank all members of Matsuishi Lab. and Hayakawa Lab. for their helpful support. Genetic specimens of whales were collected in the project of Stranding Network Hokkaido (SNH). In the specimen collection, Dr Matsuda Ayaka and Dr Kuroda Mika made a special effort to support the field collection of genetic samples used in this study. Nomurasuisan Co., Ltd., Usujirisuisan Co., Ltd. and the Usujiri Fisheries Station of Hokkaido University cooperated with the survey.

